# Is sensor space analysis good enough? Spatial patterns as a tool for assessing spatial mixing of EEG/MEG rhythms

**DOI:** 10.1101/2021.09.11.459914

**Authors:** Natalie Schaworonkow, Vadim V. Nikulin

## Abstract

Analyzing non-invasive recordings of electroencephalography (EEG) and magnetoencephalography (MEG) directly in sensor space, using the signal from individual sensors, is a convenient and standard way of working with this type of data. However, volume conduction introduces considerable challenges for sensor space analysis. While the general idea of signal mixing due to volume conduction in EEG/MEG is recognized, the implications have not yet been clearly exemplified. Here, we illustrate how different types of activity overlap on the level of individual sensors. We show spatial mixing in the context of alpha rhythms, which are known to have generators in different areas of the brain. Using simulations with a realistic 3D head model and lead field and data analysis of a large resting-state EEG dataset, we show that electrode signals can be differentially affected by spatial mixing by computing a sensor complexity measure. While prominent occipital alpha rhythms result in less heterogeneous spatial mixing on posterior electrodes, central electrodes show a diversity of rhythms present. This makes the individual contributions, such as the sensorimotor mu-rhythm and temporal alpha rhythms, hard to disentangle from the dominant occipital alpha. Additionally, we show how strong occipital rhythms rhythms can contribute the majority of activity to frontal channels, potentially compromising analyses that are solely conducted in sensor space. We also outline specific consequences of signal mixing for frequently used assessment of power, power ratios and connectivity profiles in basic research and for neurofeedback application. With this work, we hope to illustrate the effects of volume conduction in a concrete way, such that the provided practical illustrations may be of use to EEG researchers to in order to evaluate whether sensor space is an appropriate choice for their topic of investigation.

## 1. Introduction

Alpha rhythms (8–13 Hz) are a prominent feature of human non-invasive electrophysiological recordings. Different types of rhythms are found within this band, with generators in occipital, parietal, temporal and sensorimotor cortices [1, 2]. The different alpha rhythms show a functional specificity, with event-related desynchronization due to motor action for the sensorimotor rhythm, or strong modulation due to eye-opening or closing for the occipital alpha rhythm. Within each rhythm type there may be an even finer degree of organization, with differential modulation of the sensorimotor mu rhythms by hand vs. foot movements [3] or differential modulation of occipital alpha rhythms by stimuli in different parts of the visual field [4, 5]. In addition, alpha rhythms have been shown to be associated with attention showing stronger amplitude in cortical areas where neuronal activity should be suppressed [6]. In general, these rhythms remain a topic of active research directed at elucidating their role in cognition, perception and motor systems.

A fundamental challenge in the analysis and interpretation of signals recorded with electroencephalography (EEG) or magnetoencephalography (MEG) is volume conduction [7]. Volume conduction leads to overlap of signals from different generators in space and time [8]. This overlap is especially problematic for sensor space analysis, in which signals from sensors are used directly, by aid of a standard reference, e.g., common average, linked mastoids or nose-reference. Yet, despite distortions introduced by volume conduction, sensor space analysis remains a popular approach for the analysis of EEG/MEG signals [9]. The wide-spread use of sensor space analysis is certainly due to the convenience of the procedure. In contrast to sensor space, source analysis requires: 1) data analysis training in inverse modelling and understanding of its parameters, 2) more computational resources required by inverse modelling algorithms 3) more training in statistical analysis, as corrections for multiple comparisons across sources are required 4) possibly more resources need to be spent on the acquisition of individual anatomical magnetic resonance imaging data. However, despite the relative ease with which sensor space analysis can be performed, it may potentially obfuscate any fine degree of spatial specificity of neuronal rhythms to behavior. Therefore, it is of interest to assess in more detail how analysis in sensor space may blur contributions of different types of rhythms.

The methodological validity of measures derived from sensor space data is especially relevant for studies involving EEG recordings with a small number of electrodes. For instance, in a clinical setting, time constraints often limit the number of electrodes which can be placed on a patient. For instance, [10] used 1-electrode EEG to study a large cohort of patients with schizophrenia. In neurofeedback studies, typically participants receive feedback in the form of oscillatory power of a single/limited number of sensors. In closed-loop EEG studies [11, 12], where magnetic stimulation is given dependent on features of EEG rhythms, only a small number of EEG electrodes is used for the extraction of features of interest to be robust against experimental noise. If only a small number of sensors is to be used, the sensitivity of measures for this specific recording setup has to be considered in order to reliably detect the phenomena of interest.

In this article, we illustrate the impact of spatial mixing on neuronal rhythms on the sensor space level compared to the source-level. A number of studies has evaluated consistency and sensitivity of measures in sensor vs source space in the realm of connectivity metrics with respect to volume conduction and linear mixing [13, 14]. But here we focus on univariate properties of neuronal rhythms, mainly band-power of rhythms in the alpha-band. While many previous studies acknowledge the problem of volume conduction for the EEG/MEG analysis in sensor space in general, to the best of our knowledge there are no reports directly showing how individual components/sources are actually mixed at the level of sensors. We do so in this paper using specifically alpha rhythms, while the main conclusions can be generalized to other oscillations and evoked responses.

The main contribution of the following article is the quantification of spatial mixing of rhythms on the sensor space level. First, we discuss an easy-to-use method for assessing origin and spatial spread of extracted rhythms given a standard sensor scheme via the calculation of spatial patterns and demonstrate practical applications. We then use spatial patterns to assess spatial mixing of neuronal rhythms on the sensor space level compared to source level by using simulations in a realistic head model and a large dataset of EEG resting-state rhythms. Here, we illustrate constituent band-power contributions of different rhythms in the alpha-band in single sensors. Additionally, we show how spatial mixing is even more problematic when using ratio-measures of oscillations, due to the dynamic nature of oscillations, with high varying amplitude modulation of neuronal rhythms, affecting relative contributions of specific rhythms. We hope that our illustrations provide intuitions for basic and clinical researchers, in order to evaluate whether sensor space analysis may or may not be appropriate for their use case.

## 2. Materials and Methods

The analysis was performed using python and MNE version 0.23 [15] for the empirical analysis. The analysis code needed to reproduce the analysis and figures is available here: https://github.com/nschawor/eegleadfield-mixing. While we show examples for single participants in the following, it is possible to generate these types of plots for all other participants with the provided code.

### 2.1. Experimental recordings

For the empirical data analysis, we analyzed EEG data which was previously collected in the project “Leipzig Cohort for Mind-Body-Emotion Interactions” (LEMON). We summarize participant details and EEG data acquisition briefly in the following. A more extensive description of the dataset of all study components can be found in the original publication [16].

#### 2.1.1. Participants

EEG data was collected from 216 volunteers who did not have a history of neurological disease or usage of drugs that target the central nervous system. The study protocol was approved by the ethics committee at the medical faculty at the University of Leipzig (reference number 154/13-ff) and conformed to the Declaration of Helsinki. Written informed consent was obtained from all participants prior to the experiment. Data from 13 participants were excluded because the files lacked event information, had a different sampling rate, mismatched header files or insufficient data quality. In addition, the data from 4 participants was excluded because it had a low signal-to-noise ratio in the alpha-band as indicated by a 1/f-corrected spectral peak in the alpha-band below 5 dB (see Spectral analysis section for exact procedure). This resulted in datasets from 199 participants (127 male, 72 female, age range: 20–77 years)

#### 2.1.2. EEG recording setup

Scalp EEG was recorded from a 62-channel active electrode cap (ActiCAP, Brain Products GmbH, Germany). In this configuration, 61 electrodes were in the international 10-20 system arrangement, and one additional electrode below the right eye was used to monitor vertical eye movements. The reference electrode was located at FCz, and the ground electrode at the sternum. The impedance for all electrodes was kept below 5 kΩ. Data was acquired with a BrainAmp MR plus amplifier (Brain Products GmbH, Germany) at an amplitude resolution of 0.1 *µ*V with an online band-pass filter between 0.015 Hz and 1 kHz and with a sample rate of 2500 Hz. Recordings were made in a sound-attenuated EEG booth.

In the experimental session, a total of 16 blocks were recorded, each lasting 60 seconds. Two conditions were interleaved, eyes closed and eyes open, starting in the eyes closed condition. During eyes open blocks, participants were instructed to fixate on a digital fixation cross. Changes between conditions were announced with the software Presentation (v16.5, Neurobehavioral Systems Inc., USA).

### 2.2. Data analysis

#### 2.2.1. Preprocessing

We used the available preprocessed data of the LEMON dataset, with the preprocessing as applied by the data creators. The preprocessing is described briefly in the following: Raw data was downsampled from 2500 Hz to 250 Hz and band-pass filtered in the frequency range 1–45 Hz with a Butterworth filter, with filter order 4. Raw activity traces were visually inspected and outlier electrodes with frequency shifts in voltage and of poor signal quality were excluded. Data was inspected for intervals with extreme peak-to-peak deflections and large bursts of highfrequency activity and these intervals were discarded. In order to reduce the dimensionality of EEG signals, principal component analysis was used to keep principal components that explain 95% of the total data variance. Next, independent component analysis based on the Extended Infomax algorithm was performed (step size: 0.00065/log(number of electrodes), annealing policy: weight change > 0.000001, learning rate is multiplied by 0.98, stopping criterion: maximum number of iterations 512 or weight change < 0.000001). Any component that reflected eye movements, eye blinks, or heartbeat related activity was removed. The remaining independent components (mean number: 21.4, range: 14–28) were projected back to sensor space.

#### 2.2.2. Spectral analysis

As the focus here is oscillatory activity in the alpha frequencyband, we included only participants which exceeded a signal-to-noise ratio in the alpha frequency-band. For this, we used a criterion of > 5 dB as in our previous work [17]. To determine the signal-to-noise ratio in the alpha band, the frequency spectrum was computed with Welch’s method (Hann window, 1 second window length, 50% overlap). To subtract the 1/f-contribution from the spectrum, we used spectral parametrization [18]. Participants were included if at least one electrode on the midline displayed an oscillatory peak > 5 dB in the alpha band, as evaluated over the whole recording length.

#### 2.2.3. Extraction of neuronal sources

We used spatio-spectral decomposition (SSD) [19] which is a well-validated technique allowing us to extract neuronal oscillations with the maximized signal-to-noise ratio in a specified frequency band. The method is based on generalized eigenvalue decomposition of covariance matrices across sensors and maximizes the oscillatory power of a component at a specified target frequency band, while simultaneously minimizing the power at flanking frequency bands, yielding oscillatory components with highest signal-to-noise ratio. The computation can be performed fast and with few parameters. For our use case, we defined the frequency band of interest as the participant-individual peak in the alpha band, with a bandwidth of ± 2 Hz.

#### 2.2.4. Assessing spatial mixing with the aid of spatial patterns

To examine how different components mix on a chosen sensor, we analyzed the spatial pattern coefficients associated with the SSD spatial filters. The general pipeline is shown in Fig. 1. Spatial patterns describe the contribution of sources **S** on the activity recorded from sensors **X** in a linear way: **X** = **AS**, with **A** being the matrix of spatial patterns, sometimes also called mixing matrix. In our convention, the columns of the matrix contain the spatial patterns for the individual sources, and the rows of the matrix contain the contributions of the contribution of the individual sensors to each source. Spatial patterns were computed according to [20] on the basis of the covariance of activity filtered in the alpha band multiplied with the spatial filter obtained with SSD. As generalized eigenvalue decomposition methods are polarity invariant, we analyzed the absolute value of the spatial patterns in some cases.

**Figure 1:**
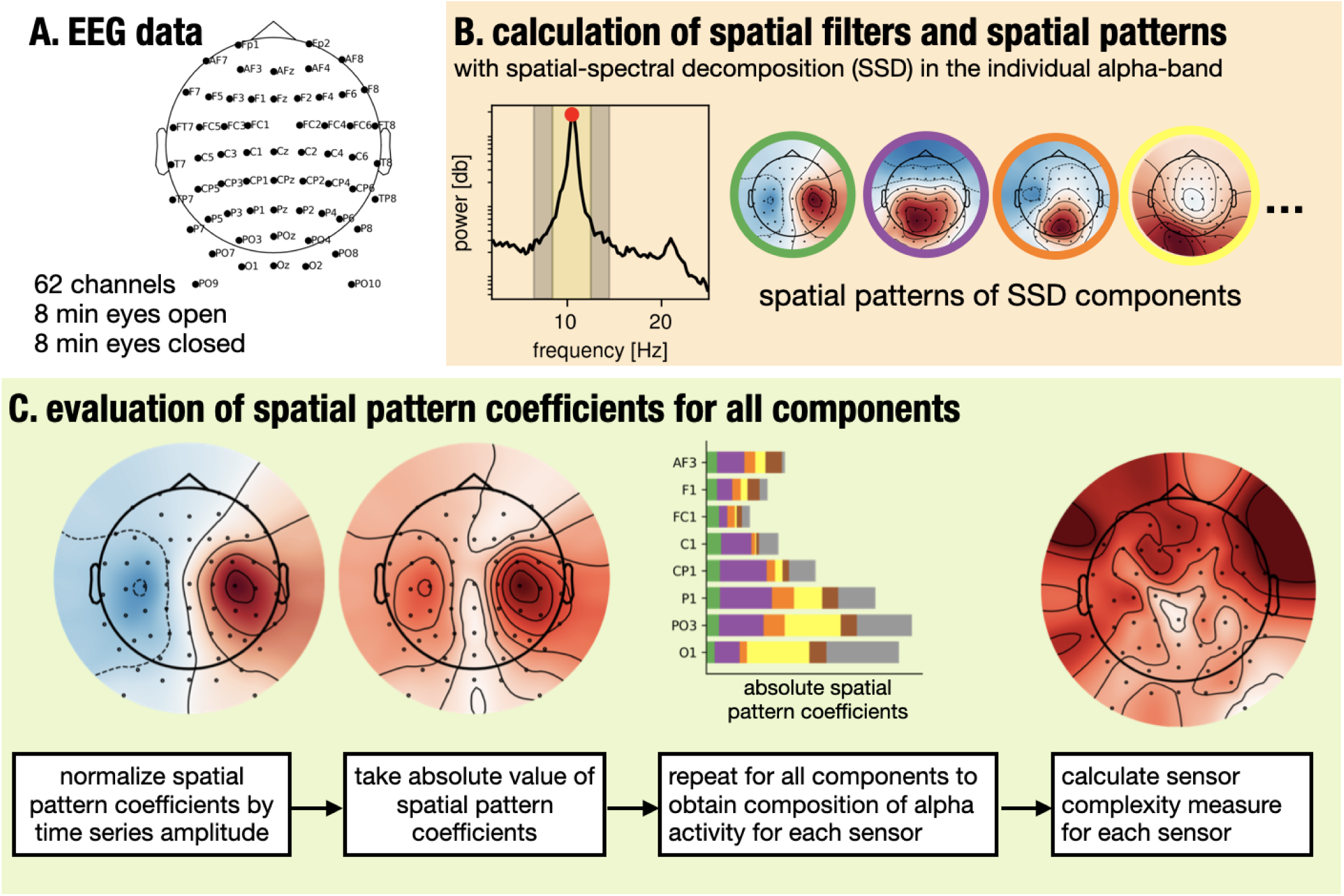
Analysis pipeline for quantifying the contributions of independent rhythms on sensor activity. A. The dataset consisted of 62-channel resting-state EEG recordings for eyes open and eyes closed conditions. B. Spatial filters and patterns were calculated with spatio-spectral decomposition (SSD) using narrow-band data in the individual spectral peak in the alpha frequency-band. C. The entries of the spatial patterns for each sensor were extracted and normalized, the absolute value was taken to calculate the sensor complexity for each EEG electrode.

To assess how rhythms contribute to each sensor, we then computed a measure quantifying the deviation from a scenario where all components contribute with equal power to the signal of a given sensor. This measure is called sensor complexity in the following and allowed us to assess the relative contribution of each source in the observed EEG activity:

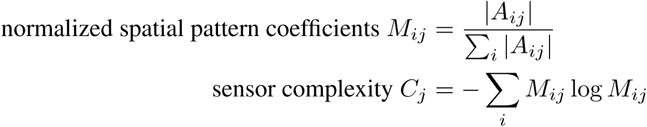

with *A*_*ij*_ as the spatial pattern coefficient for EEG electrode *j* and SSD component *i*. In the case of simulations, this is the lead field entry for a specific EEG electrode *j* and a specific source *i*. A free parameter in this context is how many components per participant are considered. Because not all components returned by SSD contain pronounced oscillatory activity in the alpha-band, we restricted the number of components to a fixed number of 10. The number of components influences the absolute value of the sensor complexity. The number of components was chosen empirically on the basis of the percentage of explained variance in the alpha-band.

### 2.3. Simulations

For the simulations, we distributed several sources of rhythms in the alpha-band in specified cortical locations in a realistic 3D head model. We then extracted the lead field coefficients for each EEG electrode and computed a sensor complexity for each sensor, which enables us to investigate spatial mixing of rhythms per sensor basis.

#### 2.3.1. Head and lead field model

We used the New York Head, a realistic precomputed lead field model of Huang et al. [21] and Haufe et al. [22]. Here we give a brief description of the generation of the head model and lead field, with full details given in the above articles. Briefly, the anatomical basis for this model is the detailed ICBM152 head model, based on the average of 152 adult brains, imaged with magnetic resonance imaging [23]. For this head model, the finite element lead field solution is provided for a set of 231 standardized electrode positions and 75,000 nodes distributed on a cortical surface mesh. We extract the lead field entries where dipole orientations are assumed to be perpendicular to the cortical surface. The New York head lead field is provided for a common average reference. For the demonstration in Fig. 4, the ‘fsaverage’ example data and head model provided by MNE was used.

#### 2.3.2. Placement of alpha generators in a 3D cortex model

Sixteen sources were placed in each hemisphere with locations approximated according to [2]. We considered six occipital, two inferior parietal, three somatosensory and five temporal alpha sources. As physiological rhythms are known to have different amplitudes, e.g., the more pronounced visual alpha rhythm, the different rhythm types were multiplied with a specified gain factor, as listed in Table 1, with higher power for occipital, parietal and sensorimotor sources and lower power for temporal sources. Additionally, we modelled a state change from eyes open to eyes closed state, during which the sources placed in the occipital region increase in strength, while other sources remain unchanged. The lead field coefficients were multiplied with the type specific gain factors for the respective conditions. The lead field entries were calculated for each sensor and visualized as a proportion on a topographic map. The original head model contains 231 EEG electrodes, the number of visualized electrodes was reduced to match the number of electrodes in the empirical data.

**Table 1:**
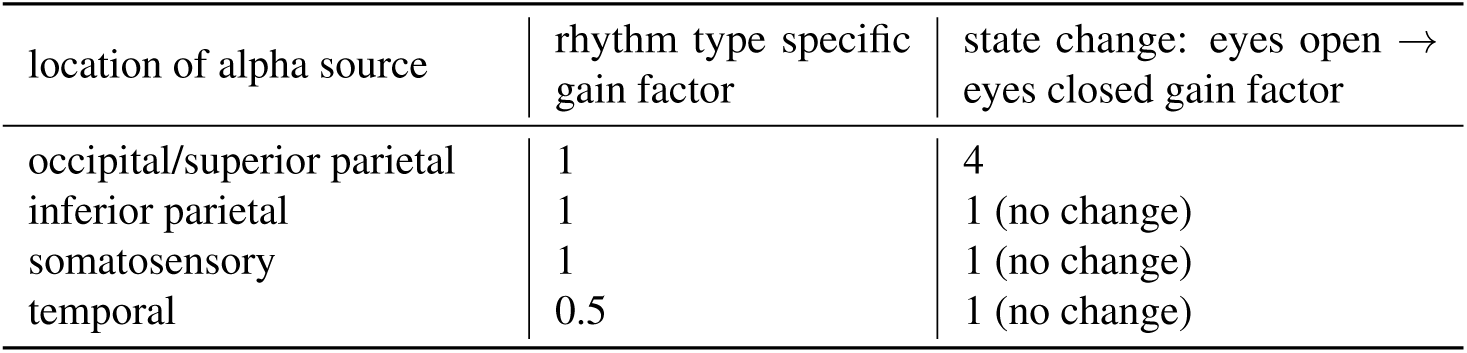
Gain factors for types of alpha activity sources, indicating their relative strength and state change properties.

#### 2.3.3. Assessing spatial mixing with the aid of the lead field

To examine how different oscillatory sources mix on a given sensor, we analyzed the lead field coefficients for each sensor. The lead field for a constrained dipole orientation is given by a matrix **L** with dimensions number of dipoles times number of sensors. Because the only sources contributing to activity in our simulations are the 16 above listed for each hemisphere, all other rows of the lead field matrix can be disregarded, resulting in 32 times number of electrodes lead field coefficients to consider. The complexity measure was calculated using the same formula as for the empirical data using the lead field coefficients weighted by the respective gain factors.

## 3. Results

### 3.1. Spatial patterns as a tool to investigate spatial correlations

First, we discuss the concept of spatial patterns. Spatial patterns are an easy way to assess the spatial distribution of activity associated with the signal from one particular sensor or spatial filter by looking at the correlation across sensors. In EEG/MEG analysis, neighboring sensors will always be correlated to a large extent due to volume conduction. Spatial patterns show how neuronal activation of sources/components in the brain maps onto EEG/MEG sensors.

In order to compute a spatial pattern, first a spatial filter needs to be defined. A spatial filter is a vector with as many entries as sensors, with a numerical weight value for each sensor. Each sensor has a certain weight in a spatial filter vector, these weights can be zero as well. For instance, the spatial filter vector for a sensor that is taken as is from the recording file without rereferencing would have an entry of 1 for that respective sensor and 0 otherwise. Referencing can be seen as the matrix multiplication of a spatial filter with the data, which yields an activity trace. Similarly, for common average referencing and Laplacian referencing a spatial filter vector can be easily constructed (see Fig. 2A). Spatial patterns are distinct from scalp potential maps, as spatial patterns reflect the spatial spread of activity originating from a specified spatial filter vector, so in the simplest case from a single sensor, whereas scalp potential maps reflect the superposition of all activity at a particular time point.

**Figure 2:**
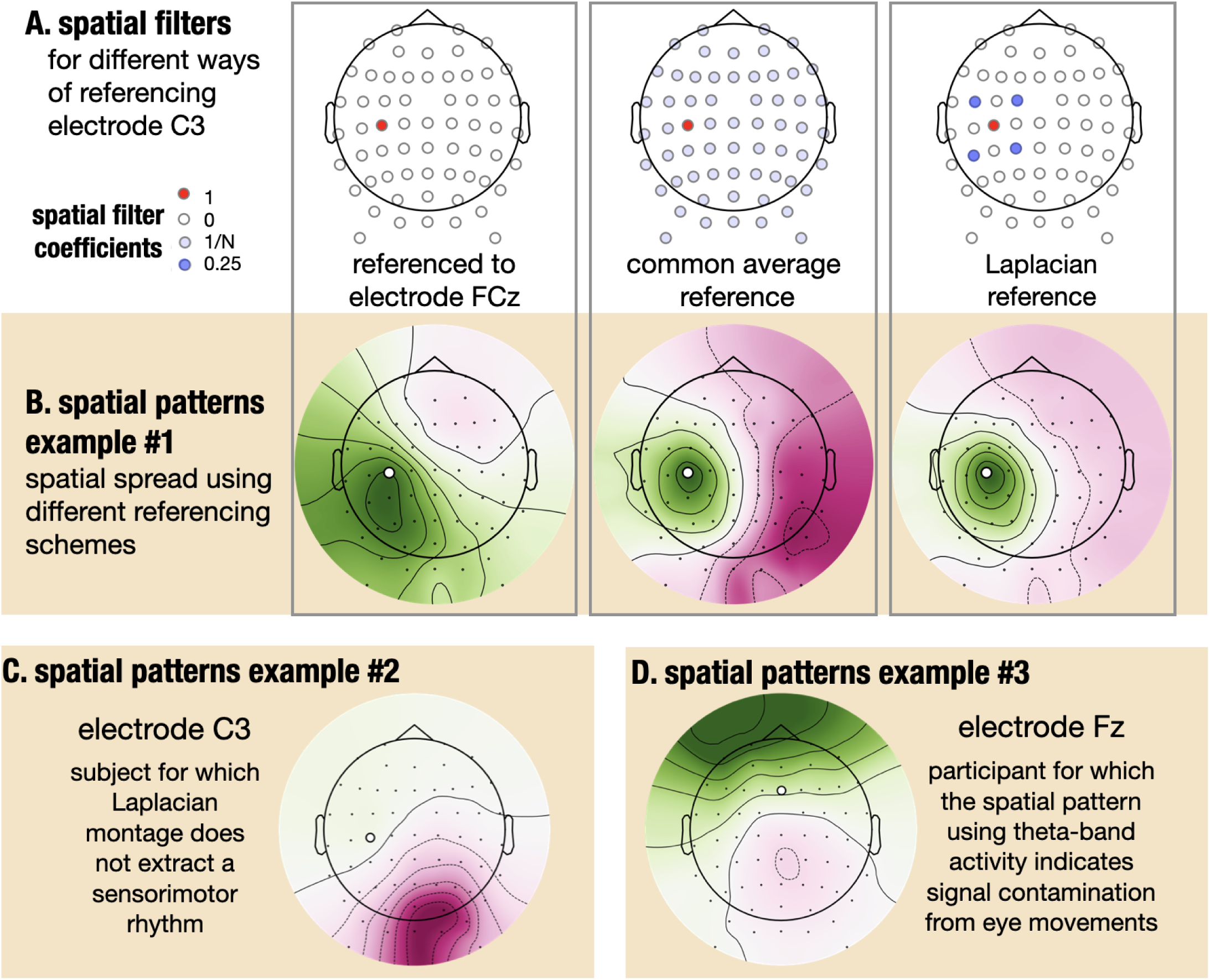
Spatial patterns aid in assessing focality and origin of extracted sensor signals. A. Spatial filters for three different referencing scenarios: referenced to electrode FCz (reference at the time of signal acquisition), common-average reference, with filter weights = 1/N with N being the number of sensors, Laplacian referenced. B. Demonstration of how activity spread is attenuated by different referencing schemes. Reference types from left to right as in A. Activity extracted with a Laplacian filter around electrode C3 shows a reduced spatial spread around the region of interest compared to referencing to electrode FCz or common average referencing. C. Demonstration of how even a Laplacian does not extract activity below the activity center, the occipital alpha activity in this participant is so strong that occipital activity shows up in the Laplacian referenced electrode C3. D. Demonstration how theta activity shows a topography reminiscent of eye movement type activity, instead of more mid-frontal distribution. because of insufficient data cleaning.

Spatial patterns are then computed by a multiplication of a specific spatial filter vector with the covariance matrix of activity across sensors. In this process, the covariance entries are added according to the polarity and strength of the spatial filter weights. The spatial filter would be equal to the spatial pattern, if the activity of sensors would be uncorrelated and the covariance matrix would be an identity matrix. But this is never the case for EEG/MEG data, therefore we need to transform spatial filters into spatial patterns in order to make statements about the location of extracted signals. For instance, the spatial pattern for a non re-referenced sensor (using the referencing at the time of data acquisition) would be exactly the covariance of this sensor to other sensors, reflecting the signal spread across sensors. The signal activity is typically band-pass filtered before computing the covariance matrix to investigate the correlation structure of the signals for a specific frequency band of interest. Different constraints can be used to calculate spatial patterns, for instance when enforcing sparsity of spatial patterns is desired, a regularization term can be used [20]. In general, spatial patterns can be seen as least squares coefficients when attempting to fit the data time series using the source time series as for instance returned by SSD.

Spatial patterns can help to verify and check the location of the signal of interest, e.g., help check for appropriate presence of oscillations to improve validity of measures. In Fig. 2B, we show the spatial patterns associated with electrode C3 over the left sensorimotor cortex, that has been referenced in three different ways: using a FCz-reference (the reference at time of signal acquisition), common average reference and Laplacian-reference, for activity in the 8–13 Hz range. It can be seen that the focality of the signal changes, depending on the respective referencing. In the ideal case, the contribution from areas far away from the chosen region should be minimized, approaching a value of 0 for the spatial pattern coefficients. It can be seen that the spatial spread is relatively broad in the FCz-referenced case and becomes more focal for a Laplacian reference. Despite improved focality for Laplacian referencing in general, the signal will not have a local origin in all cases where a Laplacian reference is used. In Fig. 2C, we show an example of a participant where applying a Laplacian filter over the electrode C3 does result in a signal originating in the vicinity of the sensorimotor cortex, but has the strongest contribution from posterior activity. In the above cases the posterior alpha activity is just very strong compared to the sensorimotor mu rhythm, which is not really detectable in this particular participant. Fig. 2D shows an example where the aim was to extract theta activity in the frequency band of 4–7 Hz using a frontal sensor, but insufficient data cleaning regarding eye movement artefacts has been performed. Therefore, the extracted activity in the theta-band is contaminated by artefacts as evident from a topography reflecting eye movements.

In summary, spatial patterns may be an easy-to-use tool for data exploration for EEG analysis. Note that all these considerations presented in Fig. 2 are in general applicable for neuronal activity in different frequency ranges and therefore these examples can be generalized to other bands, i.e., rhythms in the delta-, theta-, beta- and gamma-bands.

### 3.2. Simulations: Contribution of different alpha rhythms to sensor signals

While previously we looked at spatial patterns associated with a specific component, next we illustrate how the spatial mixing of rhythms can be assessed by analyzing multiple spatial patterns. For this, we use simulations in a realistic head model. We place 16 sources into cortical locations per hemisphere, according to [2], see Fig. 3A, with corresponding lead field entries plotted in Fig. 3B. The free parameters here are the number of sources and the strength of each source relative to others.

**Figure 3:**
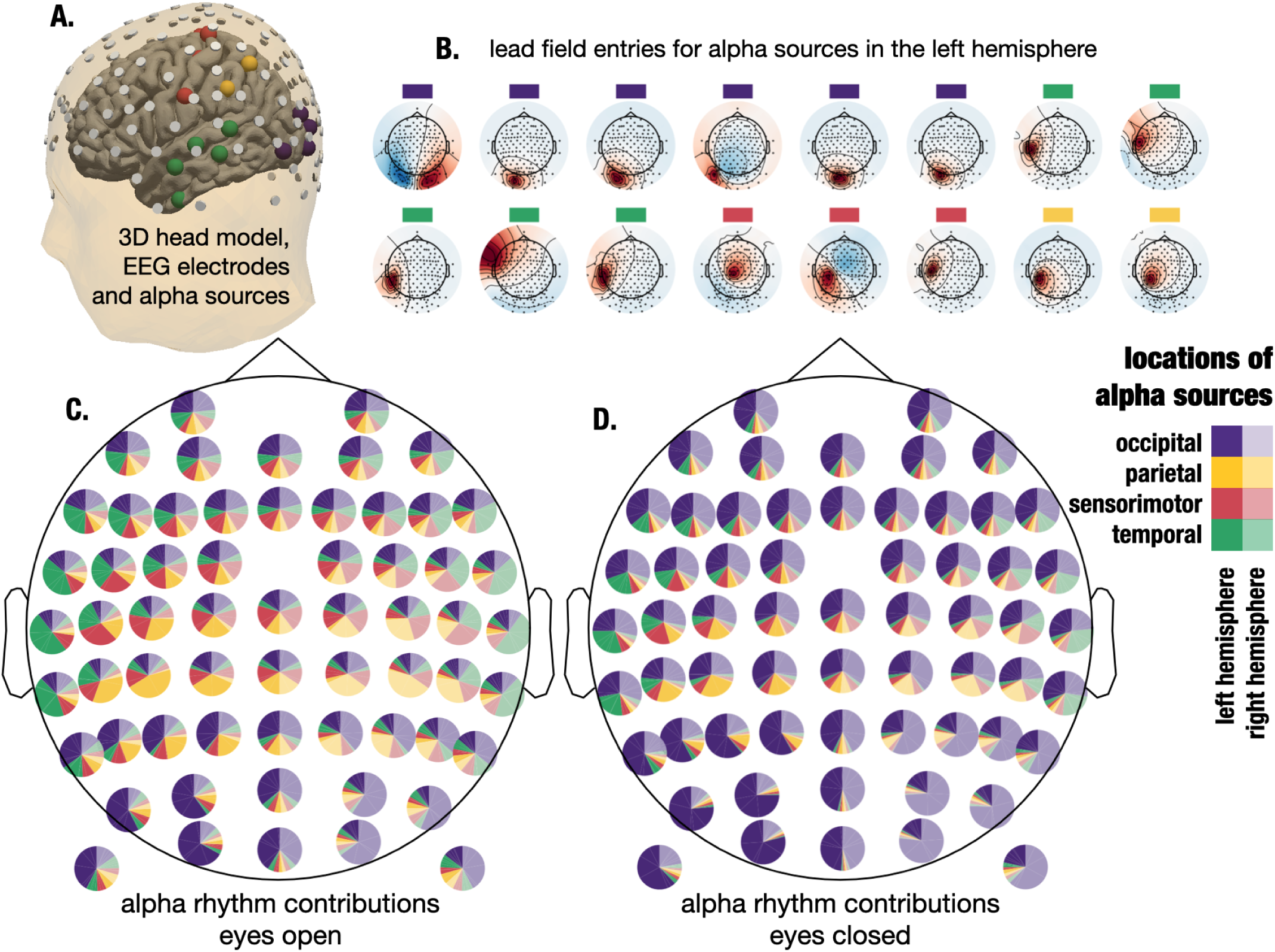
Different alpha rhythms contribute to activity recorded on each sensor, simulated example. A. 3D model of the head and cortical gray matter, with EEG electrodes and the locations of the corresponding alpha sources (blue: occipital alpha source, orange: parietal alpha source, green: temporal alpha source, red: sensorimotor mu source). B. Lead field topographies for each type of alpha source, showing contributions with positive (red) and negative (blue) polarity to the signal of each electrode for each alpha source. C. Simulated rhythm contributions onto individual sensors, eyes open condition. Each pie ploot represents one EEG electrode. The proportions displayed are colored according to rhythm type as in B, with more faint colors indicating contributions from sources located in the right hemisphere and more saturated colors indicating contributions from sources located in the left hemisphere. D. Rhythm contributions onto individual sensors, eyes closed condition, with an increased contribution of occipital alpha.

In Fig. 3C we visualize for each sensor the contribution of each rhythm by showing the absolute spatial pattern coefficient as taken from the lead field. Several observations can be noted: First, a non-trivial amount of signal is contributed from the opposite hemisphere, which may complicate the evaluation of the lateralized effects. Second, it can be seen that the majority of alpha activity at frontal sensors consists of contributions from propagated posterior alpha sources. To a large extent this is due to the orientation of the dipoles. Third, on central sensors, only a small portion of the activity in the alpha band is contributed by sensorimotor mu sources. In Fig. 3D, we show the effect of changing signal-to-noise ratio for one type of rhythm, posterior alpha. This could for instance occur in the case in an eyes-closed condition where the power of posterior alpha sources increases drastically. It can be seen that the relative contributions of visual alpha activity increase, making up a majority of the signal in the alpha band.

To further illustrate how changing the orientation of a central alpha source changes contributions in frontal sensors, we provide Fig. 4. Here, we display the location and three different possible dipole orientations in Fig. 4A with the corresponding lead field topographies in Fig. 4B and the absolute lead field coefficients for each dipole orientation in Fig. 4C and 4D. It can be seen that, while for a radial orientation of the dipole, the contribution on frontal sensors is minimal, the contribution increases for tangential orientations of the dipole.

**Figure 4:**
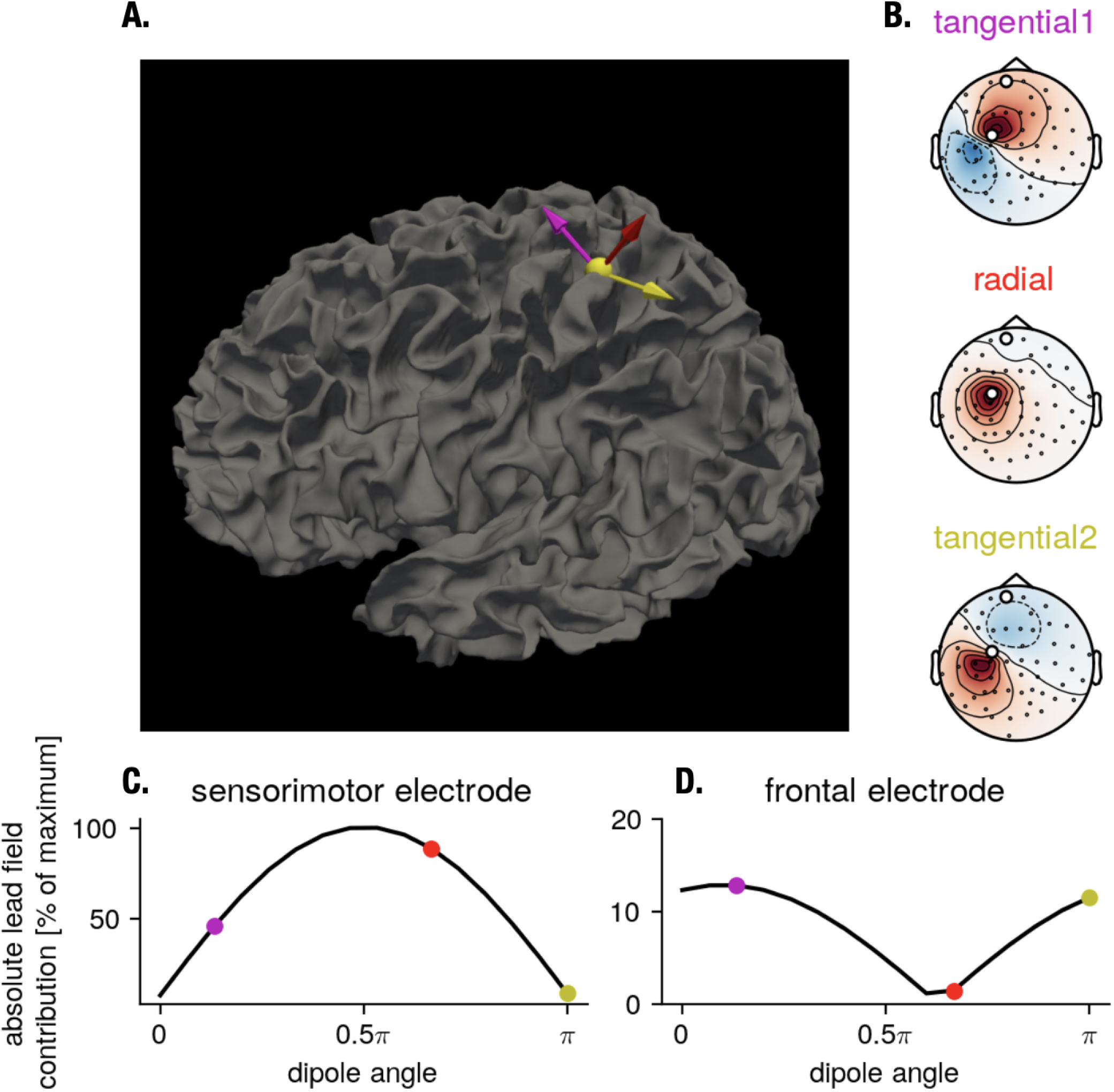
Changing the dipole orientation of a central alpha source affects sensor space activity on frontal electrodes. A. Different dipole orientations are shown on a 3D gray matter model. The color corresponds to the color in the topographies in B. B. The corresponding lead field entries for each dipole, plotted as a topography. C. Absolute lead field contribution to one sensorimotor electrode for different dipole orientations. Sensor activity is highly dependent on dipole orientation. D. Same as in C but for a frontal electrode.

### 3.3. Resting state data: Contribution of different alpha rhythms to sensor signals

To illustrate how rhythms in the alpha-band spatially overlap on sensors in empirical data, we show data for two individual participants in Fig. 5. This illustration is constructed similar to the simulation illustration shown in Fig. 3C and 3D. Since the ground truth mixing coefficients are not known for empirical data, we estimate the components and the spatial patterns using a statistical approach based on spatio-spectral decomposition (SSD). Example topographies of components are shown in Fig. 5A, ordered by signal-to-noise ratio in the alpha frequencyband. Components reflecting typical occipital alpha and sensorimotor mu rhythm topographies can be seen. In Fig. 5B, the contribution for each component onto individual sensors as evaluated in terms of band-power is shown. Fig. 5C and 5D are analog for a different participant. The figures generated are for a fixed number of components (N=10).

**Figure 5:**
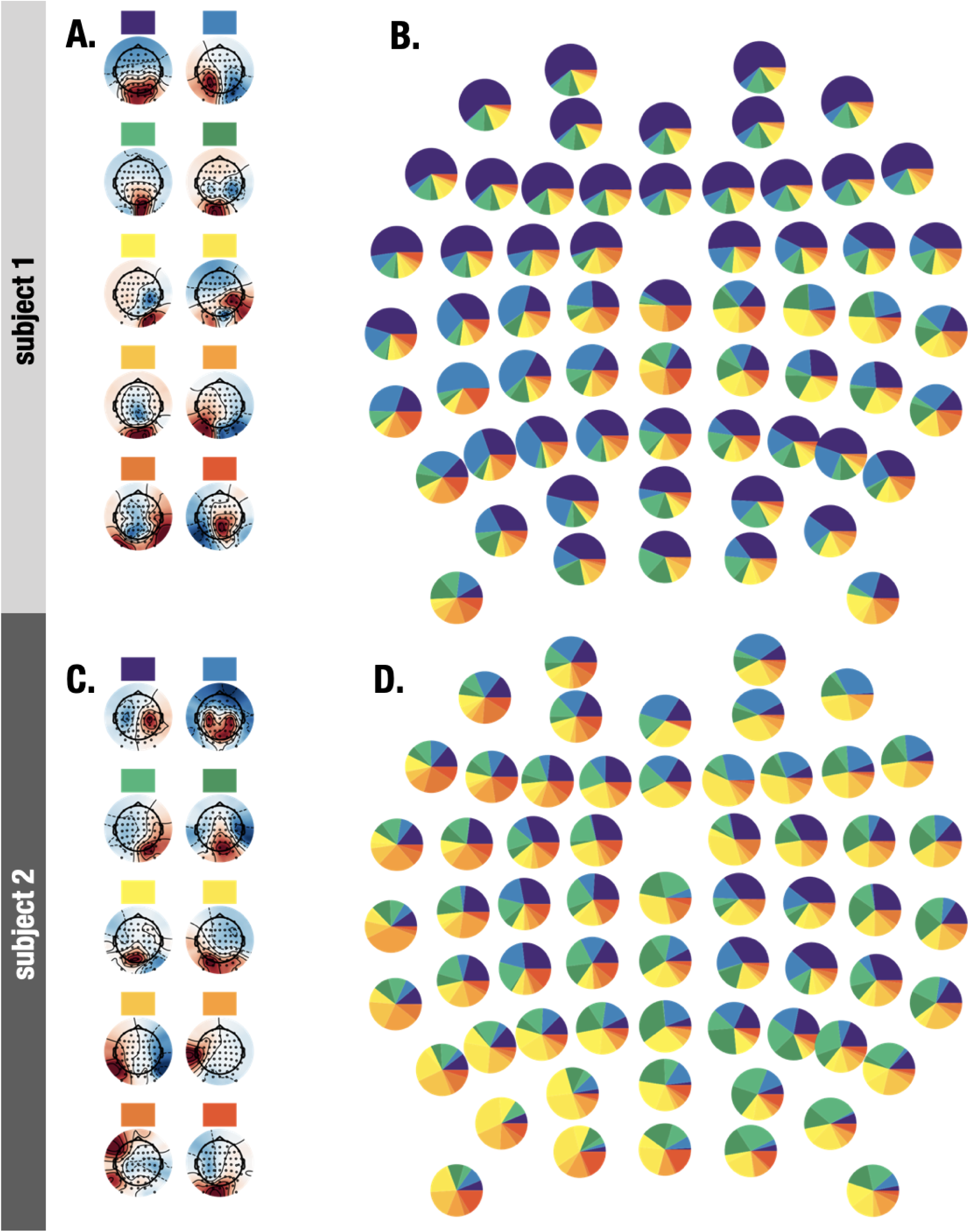
Different alpha rhythms contribute to sensor space activity, empirical example for two participants. A. The first ten patterns in the alpha-band for one participant, for the eyes open condition. Each rhythm was assigned a color which corresponds to the colors in the next subplot. B. The proportion of the ten SSD components present at each EEG electrode, as assessed with aid of the relative contribution. While sensors in the sensorimotor regions show the highest proportion of sensorimotor rhythms, also alpha rhythms originating from occipital regions contribute to the activity recorded at these sensors. C and D are analog to A and B for a different participant.

Analog to the simulation, it is evident that for frontal sensors a large part of the activity in the alpha-band is from posterior alpha components with strongest contributions to occipital and parietal sensors. Over the sensorimotor sensors, occipital alpha activity also contributes a major part to sensor space alpha activity. Since the spatial patterns are the results of an estimation procedure, the proportions may change depending on the method used for decomposition. But the overall results are in correspondence to the simulation results, hinting at the fact that some rhythms and phenomena may be easier to detect in EEG due to higher amplitude in general.

### 3.4. Resting state data: Spatial mixing across participants

After demonstrating the qualitative effect of spatial mixing in single participants, we aim to see if we can see generalities regarding spatial mixing across participants. For instance, whether we can identify sensor locations where the mixing of different rhythms is particularly pronounced and thus representing challenges for the interpretation of the electrophysiological results. We compute a sensor complexity measure for all EEG electrodes and different states (eyes open/closed) to quantify the degree of spatial mixing.

Fig. 6A and Fig. 6B show the mean sensor complexity for both eyes closed and eyes open conditions. The eyes closed condition features a much higher power for occipital alpha sources, and a large deviation from uniform contribution for occipito-parietal sensors. This is expected since only a few sources contribute a large proportion of the power in the alpha-band. For the central sensorimotor sensors, there is a relatively high complexity since here, there are contributions from the sensorimotor mu rhythm as well as from the occipital alpha rhythms. In the eyes open condition, the situation changes, since the occipital alpha sources are now much weaker and we see less spatial mixing on central sensors. In addition, we also show complexity values for individual participants in Fig. 6C and 6D for an occipital and sensorimotor sensor respectively, to demonstrate high variability regarding spatial mixing across participants.

**Figure 6:**
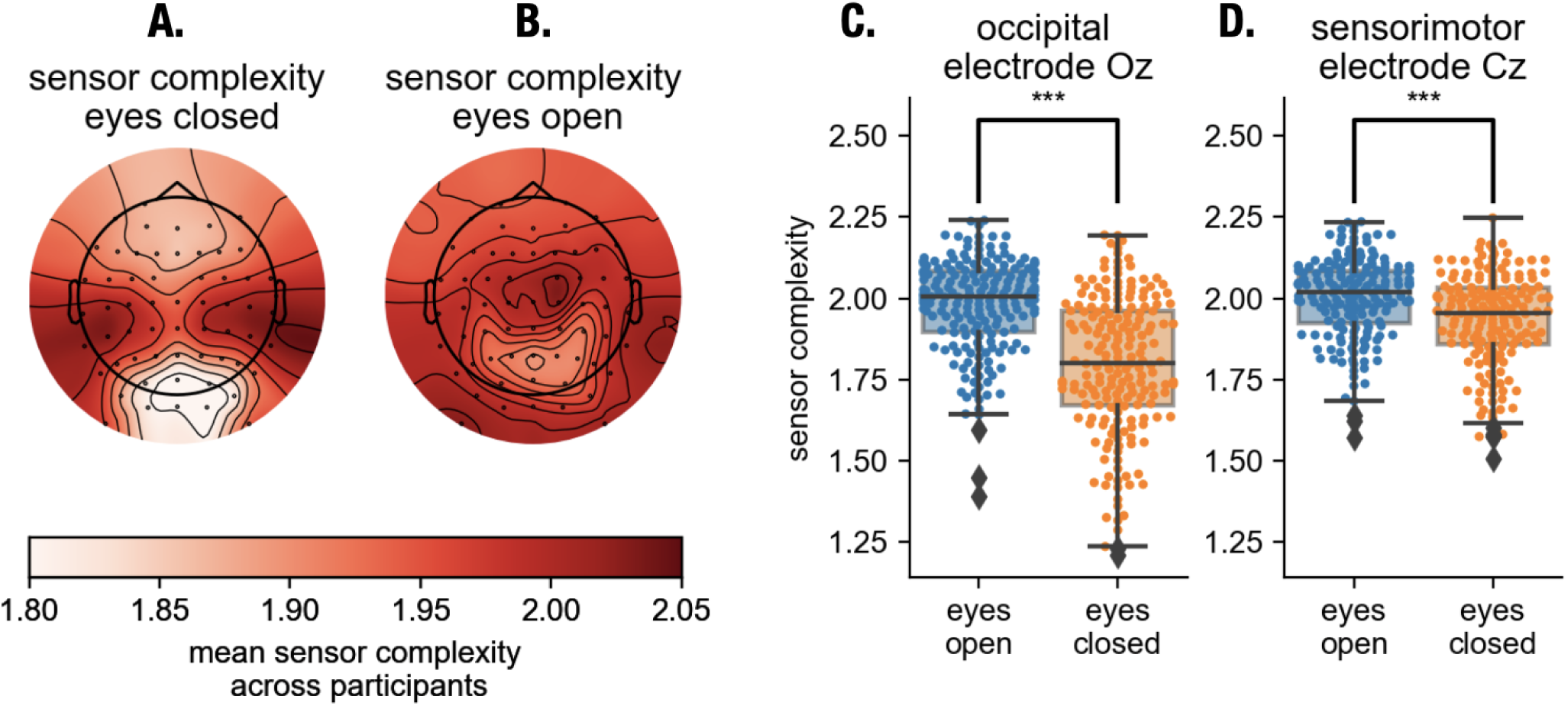
Mean sensor complexity across participants indicates less spatial mixing for posterior channels. A. Mean sensor complexity over participants for eyes open condition. B. Mean sensor complexity over participants for the eyes closed condition. Higher complexity is observed for sensorimotor sensors in the eyes closed condition, indicating a higher spatial mixing. C. Sensor complexity for individual participants for occipital electrode Oz (paired rank-sum test, p<0.0001) and D. sensorimotor electrode Cz (paired rank-sum test, p<0.0001).

### 3.5. Adding a dimension: temporal fluctuations of EEG alpha rhythms

For our calculations so far, we averaged power across time, disregarding temporal fluctuations. But neuronal oscillations also display prominent fluctuations over fast and slow time scales. Therefore, in the following we briefly illustrate oscillatory fluctuations over time for individual participants, in order to show how contributions from individual rhythms change over time for different EEG electrodes in Fig. 7A and 7B. The corresponding topographies are shown in Fig. 7C, showing sensorimotor and posterior alpha rhythms. When expressing the alpha power of SSD components as a ratio of the SSD component #2 over component #1, it can be seen the range of the power ratio between the components changing substantially over time, see Fig. 7D and 7E. Note that at different time segments the proportion/ratio of different rhythms may change. If one examines the changes in the amplitude in a frequency band of one sensor, the changes can reflect different underlying scenarios. For instance, only one source is changing or many sources are changing simultaneously. This can depend on different factors, ranging from the strength of their amplitude envelope correlations [24] or other time domain properties, e.g., whether the rhythms appear in bursts or are of more continuous nature. In general, the stronger the spatial mixing on a given sensor, the harder it is to make inferences regarding specific rhythms from the activity recorded at the specific single EEG electrode. While we show an example of one participant here, the dynamic changes of the amplitude of alpha rhythms are a general phenomenon and are present in all other participants to some extent, if they display oscillatory rhythms in the alpha-band.

**Figure 7:**
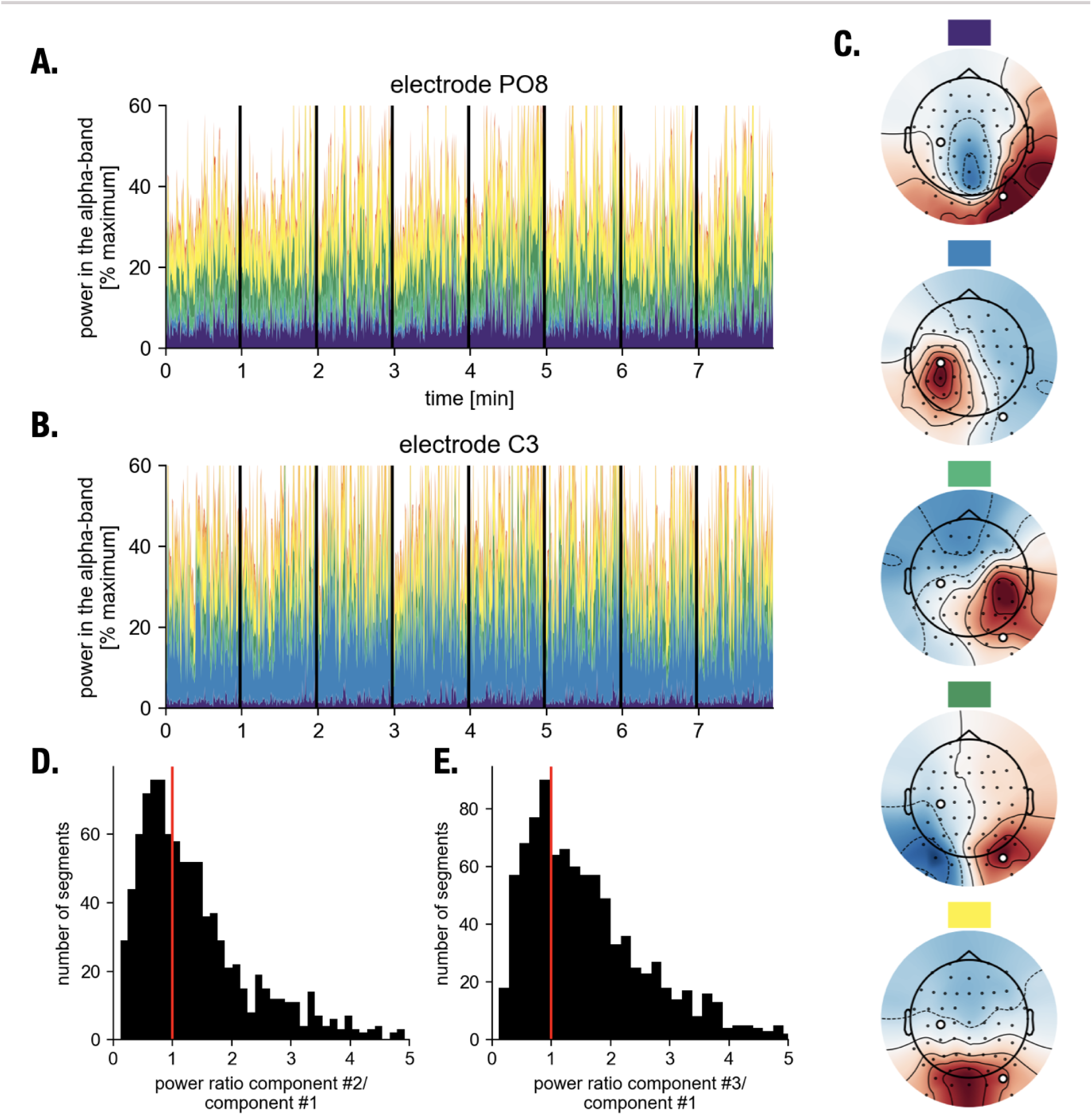
Relative alpha rhythm contributions to sensor space activity change over time. A. Time resolved contributions from different rhythms for central sensorimotor electrode C3. Vertical bars indicate block breaks. The y-axis limits are adjusted to highlight alpha power variations. B. Same as in A but for posterior electrode PO8. C. Topographies of components, color-coded as shown in A and B. E. Ratios of amplitude contributions over time for different SSD components #2 over SSD component #1 F. Same as E but for SSD component #3 over SSD component #1. Relative power contributions to sensor space activity vary substantially over time.

## 4. Discussion

With this article, we aim to raise awareness for the effects of spatial mixing on alpha rhythms as detected with EEG/MEG. We first illustrated the usage of spatial patterns to analyze focality and origin of EEG activity as a practical tool for researchers. Using this tool, we evaluated the contributions of different alpha rhythms on EEG electrodes. First, we simulated the presence of different alpha generators in a realistic head model and computed contributions using the corresponding lead field. The simulation analysis was complemented by empirical data analysis in a large dataset, where we analyzed spatial pattern coefficients for alpha rhythms as extracted by SSD. A complexity measure on individual sensor level was defined and used to illustrate how alpha sources map onto EEG electrodes, also depending on state.

### 4.1. Implications

#### 4.1.1. Amplitude of rhythms and alpha asymmetry measures

To date, many EEG/MEG studies are performed in sensor space. One of the clear advantages of such an approach is its relative technical simplicity not requiring source analysis using biophysical or statistical constraints (for example using independent component analysis or SSD). Typical examples include spectral analysis, amplitude dynamics, e.g., event-related desynchronization/synchronization [25], microstates [26], diverse complexity measures such as long-range temporal correlations [27], approximate and sample entropy [28]. A typical approach in such studies is to define regions of interest on the basis of spatial locations of sensors, for instance frontal, central temporal, parietal and occipital regions. This is often done with the hope that the activity picked-up by the sensors in these regions of interest would reflect cortical processes generated in the proximity of these sensors. However, as one can see from the simulation illustrated in Fig. 3A very large part of activity detected in frontal sensors can originate from the occipital sources. This situation is particularly important for the inference regarding alpha sources calculated on the basis of sensor space activity in electrodes F3 and F4. The asymmetry in alpha power between these EEG electrodes is often used as an indication for making conclusions about approach/avoidance behavior [29]. In this context, a stronger activation of the left hemisphere (smaller alpha power) indicates a tendency toward approach behavior while a stronger activation of the right hemisphere indicates rather avoidance. These conclusions are naturally based on the assumption that alpha activity in these frontal electrodes reflect neuronal processing, for instance in dorsolateral prefrontal cortex. However, this assumption can be very misleading. In fact, our analysis shows that the contribution of a combination of occipital and central sources can be as high as 75% in frontal sensors. This in turn makes inferences about the activation of the dorsolateral prefrontal cortex on the basis of frontal electrode activity quite problematic. Moreover, using real data, Fig. 5 shows that many occipital and central sources contribute to the power of alpha rhythms in frontal electrodes. On the one hand, it’s possible to investigate alpha asymmetry in different pairs of electrodes to show that primarily asymmetry in the frontal electrodes corresponds best to the behavioral quantification of approach/avoidance traits. However, such conclusions would not necessarily be correct since mixing of alpha rhythms might be more complex/different in occipital areas compared to frontal ones and thus asymmetry of alpha sources outside of frontal areas can still be a major contributing factor for alpha asymmetry in frontal electrodes [30]. In general, we would recommend to perform some simple decomposition of alpha sources with independent component analysis or SSD to calculate the proportion of components with clear central and occipital patterns to the whole power at frontal electrodes. If this proportion is more than 50% a caution should be applied when interpreting frontal alpha asymmetry. Such decompositions can be performed even when the recording consists of approximately 20 EEG electrodes since spatial patterns of the components could be identifiable as having central, frontal or occipital sources.

A similar logic can be applied to other locations of electrodes and other phenomena where the power of oscillations or their asymmetry should be deduced. For instance, for the sensorimotor mu rhythm, an oscillatory power difference between two hemispheres can indicate asymmetry in excitation/inhibition-balance between the hemispheres on the basis of which a certain therapeutic transcranial magnetic stimulation protocol can be prescribed [31]. In this case, a careful evaluation of alpha-band mixing complexity is also important if one is using standard reference schemes such as those based on common average, linked mastoids etc. Again, we would like to emphasize that for a more refined spatial estimation a source analysis is preferred.

#### 4.1.2. Neurofeedback in sensor space

Another important example for the use of alpha power, obtained in sensor space, is neurofeedback. Here, the main idea is to volitionally up- or down-regulate power of oscillations at a specific sensor location [32]. The main premise is that the changes in alpha power are likely to be associated with functional changes of the corresponding neuronal networks. Typically, a relationship is assumed between the power of alpha rhythms and a spatially restricted neuronal network generating these alpha rhythms. However, our simulations show that power in a given sensor reflects activity from generators in a variety of different brain areas. Therefore, no exact correspondence between the increase of oscillations e.g., at electrode Pz and spatial activation in a given cortical patch can be established, even if activation is defined quite broadly, i.e., frontal, central or occipital locations. Moreover, the power ratio of different SSD components varies as a function of time (see Fig. 7) thus further obscuring a relationship between changes of alpha rhythms and underlying neuronal processing. Such complexity of spatial mixing should inevitably lead to a decrease in the efficacy to learn neurofeedback since reinforcing a specific power of alpha rhythms at a given sensor biologically would correspond to reinforcing undetermined and ever-changing patterns of corresponding neuronal activity. This can be one of the reasons for the observation that many participants are not able to learn neurofeedback effectively [32]. In fact, on the basis of our results we hypothesize that the participants with the lower spatial complexity of alpha rhythms should be more efficient in performing reliably in neurofeedback sessions. This can be tested directly in future studies. Since neurofeedback typically requires multiple sessions and this is a time-consuming procedure, as a practical recommendation we suggest performing at least one recording with a high number of sensors (for instance 60) in order to quantify the presence and spatial complexity of alpha rhythms at different sensors.

One can then determine sensors with sufficiently low complexity to be used later with low-electrode montages (for multisession training) or in case of participants with high spatial complexity, one can proceed with more electrodes in order to enable visualizations of spatial patterns corresponding to spatially restricted neuronal activity for validation of the paradigm.

#### 4.1.3. Spatial complexity and connectivity

Previous studies have already explored effects of volume conduction on the calculation of connectivity relationships based on coherence or phase locking values [13, 14]. Here, a spurious connectivity can be detected when the same neuronal source is mapped to many sensors and therefore a high connectivity value does not reflect functional interactions but rather the fact that the same neuronal trace is mapped to different sensors thus leading to high coherence of phase locking. Clearly, volume conduction is also the reason for complex spatial patterns obtained in the present study. While we will not describe strategies to overcome detection of spurious interactions here, as it has been done in previous studies [13, 33], we want to emphasize another important aspect relating to our findings. Sensors, reflecting a high degree of spatial mixing of different components, are also likely to reflect a rich structure of neuronal interactions which can be picked up with different graph theoretical metrics even when controlled for volume conduction. Therefore, we suggest that if connectivity studies are based on a sensor space analysis, a complementary spatial sensor complexity can be computed in order to assess the possibility of obtaining hub structures particularly in sensors with the highest sensor complexity.

### 4.2. Limitations

For the empirical data analysis sections, we used a simple method for source reconstruction. With SSD, as with any other decomposition technique, it is not possible to separate all individual alpha rhythms. After all, we only record data with 60 EEG electrodes and there are many more generators than that. Therefore, the decomposition will feature components that are not of a dipolar structure, where multiple sources that are highly co-active have been combined into a single source by the decomposition algorithm. While improvements can be made in this regard, by using more sophisticated source reconstruction algorithms, our general statement is not dependent on the specific source reconstruction method we used: the activity of a single EEG electrode will reflect multiple sources in the alpha-band, for which the contributions will dynamically vary across time. In general, the existence of statistical based source separation techniques like SSD makes investigation of rhythms in source/component space easy and allow separation of individual rhythmic contributions without anatomical head models, to best utilize information from electrophysiological data.

## 5. Conclusion

Spatial mixing due to volume conduction is inherent to data recorded with EEG/MEG. Here, we have shown the extent of spatial mixing of different alpha-type rhythms and elaborated on the consequences in terms of activity contributions to sensor space activity. For detecting relationships between EEG/MEG signatures and behavior, the signal-to-noise ratio available needs to be carefully considered. While prominent posterior rhythms show less spatial mixing in sensor space, the situation is more complicated for sensorimotor and temporal alpha rhythms of smaller amplitude, potentially compromising analyses that are solely conducted in sensor space. We hope that the provided practical illustrations may be of use to EEG researchers for evaluation whether sensor space is sufficient for their topic of investigation.

## Data Availability Statement

The EEG data was previously collected as part of the “Leipzig Cohort for Mind-Body-Emotion Interactions” dataset (LEMON). The EEG data is available at: http://fcon_1000.projects.nitrc.org/indi/retro/MPI_LEMON.html. Code underlying all figure generation and analysis is available at: https://github.com/nschawor/eeg-leadfield-mixing.

